# Variation in cell surface hydrophobicity among *Cryptococcus neoformans* strains influences interactions with amoeba

**DOI:** 10.1101/867598

**Authors:** Raghav Vij, Conor J. Crawford, Arturo Casadevall

## Abstract

*Cryptococcus neoformans* and *Cryptococcus gattii* are pathogenic fungi that cause significant morbidity and mortality. Cell surface hydrophobicity (CSH) is a biophysical parameter that influences the adhesion of fungal cells or spores to biotic and abiotic surfaces. *C. neoformans* is encased by polysaccharide capsule that is highly hydrophilic and is a critical determinant of virulence. In this study, we report large differences in the CSH of some *C. neoformans* and *C. gattii* strains. The capsular polysaccharides of *C. neoformans* strains differ in repeating motifs, and therefore vary in the number of hydroxyl groups, which along with higher-order structure of the capsule, may contribute to the variation in hydrophobicity that we observed. For *C. neoformans*, CSH correlated with phagocytosis by natural soil predator *Acanthamoeba castellani*. Furthermore, capsular binding of the protective antibody (18B7), but not the non-protective (13F1) antibody altered the CSH of *C. neoformans* strains. Variability in CSH could be an important characteristic when comparing the biological properties of cryptococcal strains.

**IMPORTANCE:** The interaction of a microbial cell with its environment is influenced by the biophysical properties of a cell. The affinity of the cell surface for water, defined by the Cell Surface Hydrophobicity (CSH), is a biophysical parameter that varied amongst different strains of *Cryptococcus neoformans*. The CSH influenced the phagocytosis of the yeast by its natural predator in the soil, Amoeba. Studying variation in biophysical properties like CSH gives us insight into the dynamic host-predator interaction, and host-pathogen interaction in a damage-response framework.

## INTRODUCTION

The encapsulated basidiomycetes that comprise of the *Cryptococcus* species complex include several pathogenic species including *C. neoformans* and *C. gattii. Cryptococcus* spp. have a worldwide geographic distribution and are unusual among fungal pathogens, in that they have polysaccharide capsules that are essential for mammalian virulence.

Human infection usually begins in the lung. Infectious propagules of *C. neoformans*, in the form of spore or yeast, may be inhaled to cause a pulmonary infection that is usually cleared in immunocompetent hosts, or becomes latent. Conditions that impair immunity, such as HIV infection, are associated with disseminated disease, which usually manifests clinically as a meningoencephalitis. Recent evidence suggests that the nature of the infectious propagule has a significant effect on the outcome of the infection, as spores from *C. neoformans* cause significantly higher fungal burden in the brain of a murine model in comparison to small encapsulated yeast (1).

*C. neoformans* have been isolated from avian guano, soil, or arboreal sources. *C. gattii* has been isolated from trees, soil, freshwater, and seawater. There are three serotypes of *C. neoformans*, now referred to as *Cryptococcus neoformans* var. *neoformans* (Serotype D), *Cryptococcus neoformans* var. grubii (Serotype A) and hybrid (Serotype AD). Phylogenetic evidence suggests that they may be classified as separate species, *C. neoformans, C. deneoformans* and hybrid, respectively (2). Interestingly, *C. neoformans* var. *grubii* has been isolated from 63% of clinical samples collected world-wide, followed by *C. neoformans* hybrid (6%), and *C. neoformans* var *neoformans* (5%) (3, 4). The genomic diversity in the Cryptococcal species complex may contribute to differences in the biophysical properties of cell surfaces within the *Cryptococcus* species complex.

*C. neoformans* and *C. gattii* cells are surrounded by a polysaccharide capsule that can dramatically vary in size during infection (5), and helps the pathogen evade the mammalian immune system. Highly branched polysaccharides (6) radiate outward form the cell wall, to form a dense matrix whose porosity increases with the distance from the cell wall (7). The capsule is primarily composed of glucuronoxylomannan (GXM, 98%), along with minor components galactoxylomannan and mannoproteins. GXM contains a core repeating structure of a α-(1→3)-mannose triad, with a β-(1→2) glucuronic acid branch on every third mannose (8). The capsule of different serotypes of *C. neoformans* and *C. gattii* have distinguishable polysaccharide motifs characterized by a varied degree of β-(1→2) or β-(1→4) xylose substitutions, and 6-O-acetyl substitutions along the mannan backbone (9). Polysaccharides are highly enriched in hydroxyl groups and form an extensive network of intramolecular and intermolecular hydrogen bonds, which includes bonding with water molecules. Therefore, polysaccharides are intrinsically hydrophilic molecules, which could provide an explanation for approximately 95% of the capsule’s weight (10). Branching and substitution of polysaccharides effects the intra- and intermolecular hydrogen bonds and rigidity of the polymer, thereby effecting the polysaccharide’s ability to form hydrogen bonds with water, which results variation in hydrophobicity (11–13).

Natural variation in biophysical parameters of the microbial surface of the *Cryptococcus* species complex has been previously described. Melanization, capsule induction, and binding of capsular antibody alter the cell surface charge, which also varies by strain (14). Chronological aging of the yeast and antibody binding alter the elasticity of the polysaccharide capsule that surrounds the *C. neoformans* cell (15, 16).

CSH is a property of a microbial surface that reflects the affinity of components of the microbe’s cell surface for water and, is calculated by estimating the affinity of cell surfaces to hydrophobic substances like hydrophobic columns, solvents, or polystyrene beads (figure 1). The biological role of the CSH has been studied in bacteria such as *Staphylococcus aureus* and some fungi, and has succinctly reviewed in (17). Previous studies of *Candida albicans* have established the importance of CSH for the interaction of the pathogen with the host tissue (18). Furthermore, strain-specific variation in CSH of clinical isolates, and variation between species of *Candida* species complex have been reported (19).

**Figure 1:**
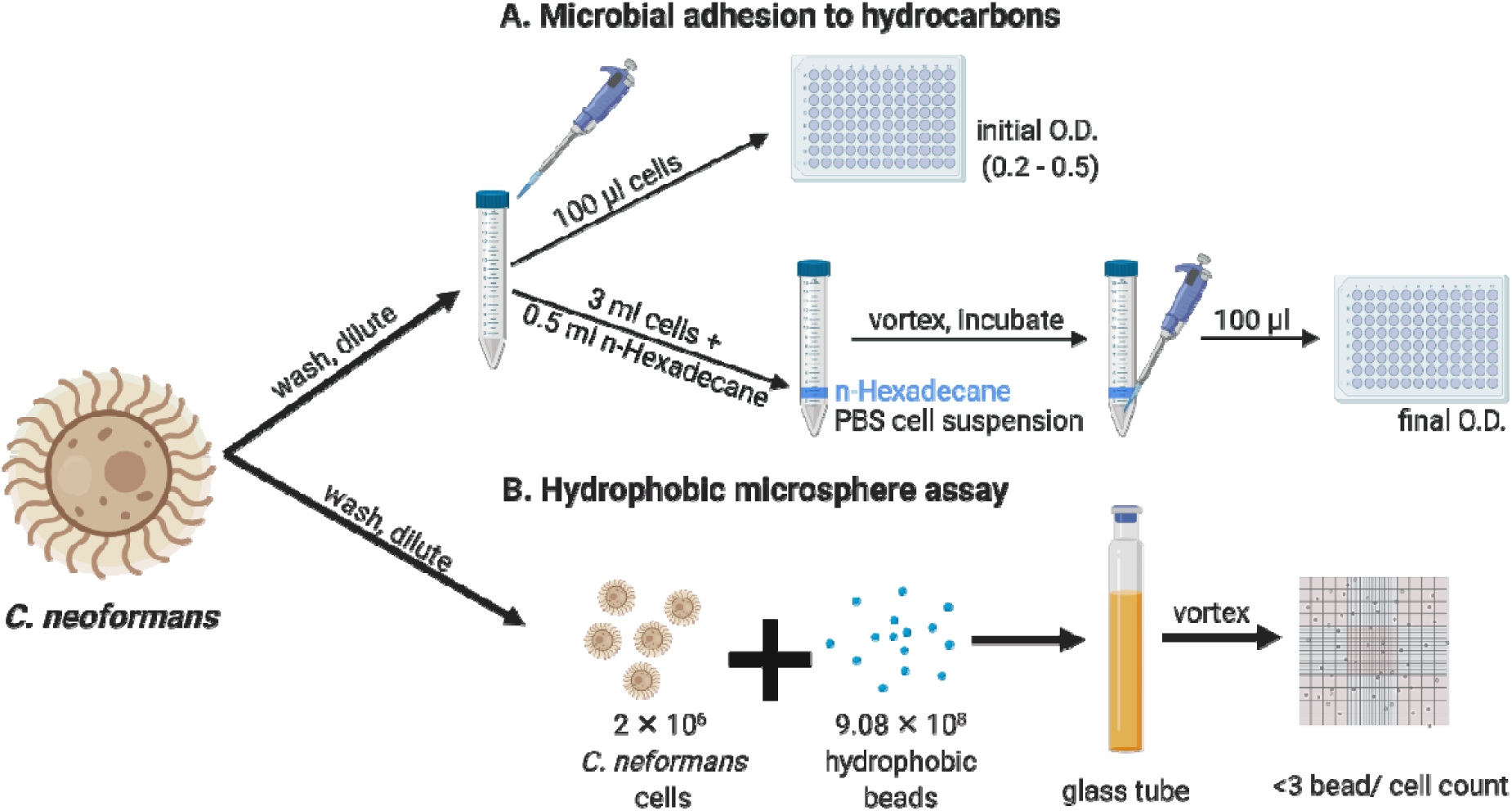
Methods for estimation of *C. neoformans* CSH. **A.** CSH estimated by MATH assay that quantifies the interaction of *C. neoformans* cells in a suspension with the hydrocarbon solvent n-Hexadecane. CSH% was calculated as the percentage change in OD of a *C. neoformans* cell suspension after vortexing the mixture of cells with n-Hexadecane. **B**. In addition, we estimated CSH by visualizing the interaction between *C. neoformans* cells and hydrophobic beads (0.8 µm) in a hemocytometer and counting cells that had >3 beads/ 100 cell to calculate CSH%. Image created with BioRender.

The biophysical properties of the infectious propagule of C. *neoformans* in the form of yeast or spore influence the interaction of the yeast with its environment, and inside the host during infection. For example, during infection, *C. neoformans* interacts with lung epithelial cells, macrophages and can pass through the blood-brain barrier. In the environment, *Cryptococcus* species complex is believed to interact with amoeba (20) and, nematodes (21). Furthermore, hydrophobicity may influence the phagocytosis of microbial cells or particles by Amoeba (22).

In this study, we report variation in CSH of *C. neoformans* and *C. gattii* strains using two independent methods. Further, we observed that CSH correlated positively with phagocytosis by *A. castellani*. Additionally, the higher order structure of the capsule is affected by the different capsular polysaccharide motifs, that vary between serotypes of *C. neoformans* and *C. gattii*, which may influence the CSH. We also found that binding of protective, but not non-protective antibodies altered the hydrophobicity of *C. neoformans* grown in capsule induction medium.

## RESULTS

### *Cryptococcal* spp. manifest significant differences in CSH

Measuring CSH by the MATH and hydrophobic microsphere techniques (figure 1) revealed considerable variability among cells of *C. neoformans* and *C. gattii* strains cultured in Sabouraud Dextrose Broth (figure 2). By MATH assay, we found that serotype D strains B3501 and JEC21 were significantly more hydrophobic than the reference strain H99 (figure 2A). By the hydrophobic microsphere assay, we found that all strains of serotype D for which CSH was estimated, including B3501, ATCC24067 and JEC21, were significantly more hydrophobic than the reference strain H99 (figure 2B). However, there was considerable strain-to-strain variation and no pattern emerged regarding differences between serotypes or species, except for the notable finding that the most strains manifesting highest CSH were *C. neoformans* serotype D.

**Figure 2:**
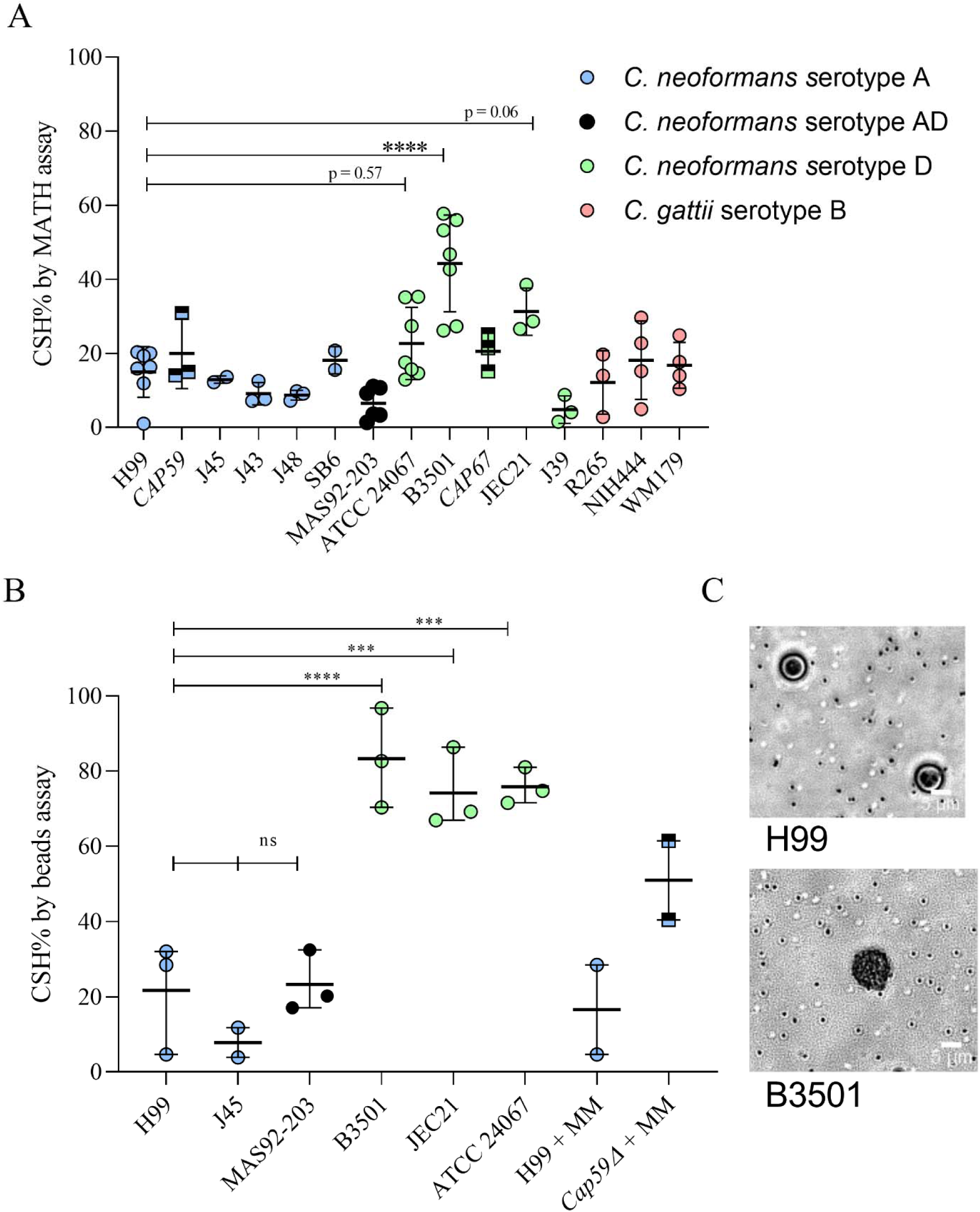
CSH of *C. neoformans* differs by strain. Graphical representation of CSH of *C. neoformans* and *C. gattii* strains. **A**. Graphical representation of CSH estimated by MATH assay. **B**. Graphical representation of CSH estimated by hydrophobic microsphere assay (left). Experiments have been performed 2-6 times independently, as indicated by individual data points. (○) Indicates a data point of CSH of an encapsulated strain of *C. neoformans* and *C. gattii*, while, (□) indicates the CSH of an acapsular mutant of the preceding *C. neoformans* strain (starting from the y-axis). Error bar represents the standard deviation about of mean. **C**. Representative image of a mixture hydrophobic beads with *C. neoformans* strain H99 (upper right) and relatively hydrophobic *C. neoformans* strain B3501 (lower right) used for the assay. Hydrophobic beads (small spheres, approximately 0.8µm in diameter) adhere to the cell surface due to the high hydrophobicity of B3501 cell, covering it almost completely. The hydrophobic beads are all but absent from the surface of H99 cells. Ordinary one-way ANOVA was used to compare the CSH of *C. neoformans* strain H99 with the CSH of *C. neoformans* and *C. gattii* strains (supplementary materials table S2). The following symbols were used to annotate the statistical significance of the results: ns, p□>□0.05; *, p□≤□0.05; **, p□≤□0.01; ***, p□≤□0.001; ****, p□≤□0.0001.

### *C. neoformans* capsule and CSH

The capsule is highly hydrophilic and is primarily composed of water (10). Hence, we sought to ascertain its contribution to CSH in *C. neoformans* strain H99 (serotype A) by comparing encapsulated H99, and non-encapsulated strain *CAP59*. To our surprise, we observed no major difference in CSH between H99 and *CAP59* cells grown in Sabouraud-dextrose broth, by the MATH assay (p = 0.9988, figure 2A, table S1). However when grown in capsule inducing minimum medium (23), the non-encapsulated strain bound more hydrophobic beads than the encapsulated strains (figure 2B). Next, we compared the CSH of *C. neoformans* strain B3501 (serotype D) to the un-encapsulated strain *CAP67*, which has a mutation in *CAP59* gene of B3501 strain (24). We observed a significant decrease in the CSH by MATH assay (p = 0.0078, figure 2A, table S1).

Different strains and serotypes of *C. neoformans* and *C. gattii* have different dominant carbohydrate motifs in their capsule (9) that my influence the experimentally observed variation in CSH. To test this hypothesis, we used *in silico* method described by Mannhold *et. al* (25), to calculate and compare the lipophilicity (log *P*) of the four dominant GXM motifs. We observed the following trend in the predicted lipophilicity of GXM carbohydrate motifs; M4 (dominant in serotype C, log *P* 2.12) > M3 (dominant in serotype B, log *P* 2.01) > M2 (dominant in serotype A, log *P* 1.9) > M1 (dominant in serotype D, log *P* 1.79).

Based on the rationale that polysaccharides enriched in greater number of hydroxyl groups would have higher hydrophilicity, we counted the number of hydroxyl groups of each dominant GXM motif (figure 3). The M4 motif (dominant in serotype C) contained the highest number of hydroxyl groups, 21, followed by 19 hydroxyl groups in M3 (dominant in serotype B), 17 hydroxyl groups in M2 (dominant in serotype A) and 15 hydroxyl groups in M1 (dominant in serotype D).

**Figure 3:**
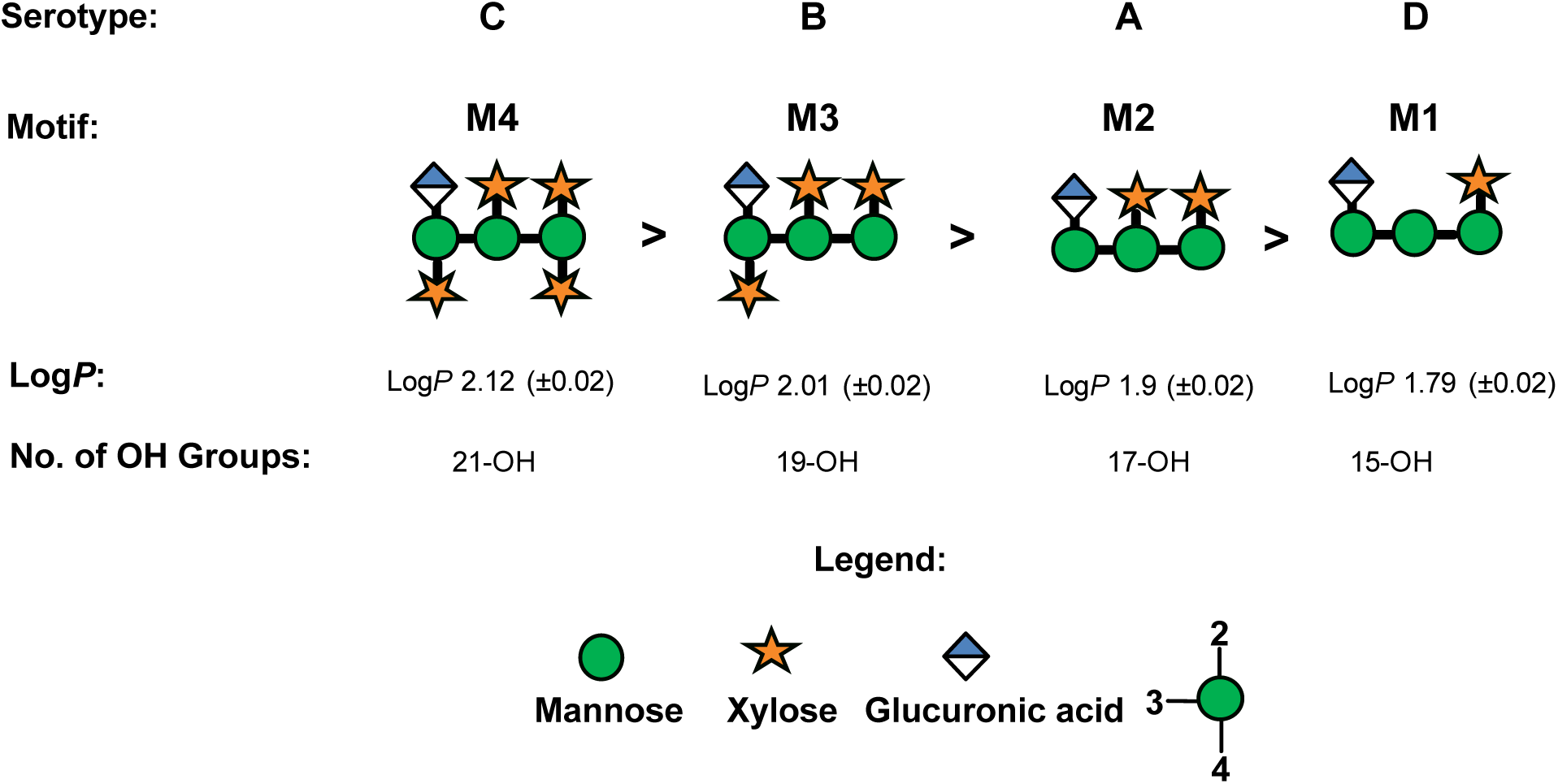
Comparison of hydrophobicity between different capsule motifs dominant in *C. neoformans* and *C. gattii*. Lipophilicity, log *P*, of dominant carbohydrate motifs in the carbohydrate was predicted by an equation proposed by Mannhold *et. al* (25). M4 was found to be the most hydrophobic motif and M1 the least. The number of hydroxyl groups on each polysaccharide motif was calculated (below). Glycan notification followed the Symbol Nomenclature for Glycans (SNFG) (67).

### CSH of unopsonized *C. neoformans* correlates with phagocytosis by *A. castellani*

To test whether CSH influences phagocytosis by soil predators like the amoeba, we incubated fungal and protozoal cells and estimated the phagocytosis index. We found a positive and linear correlation between CSH of *C. neoformans* strains and phagocytosis index of *C. neoformans* strains by *A. castellanii* (figure 4).

**Figure 4:**
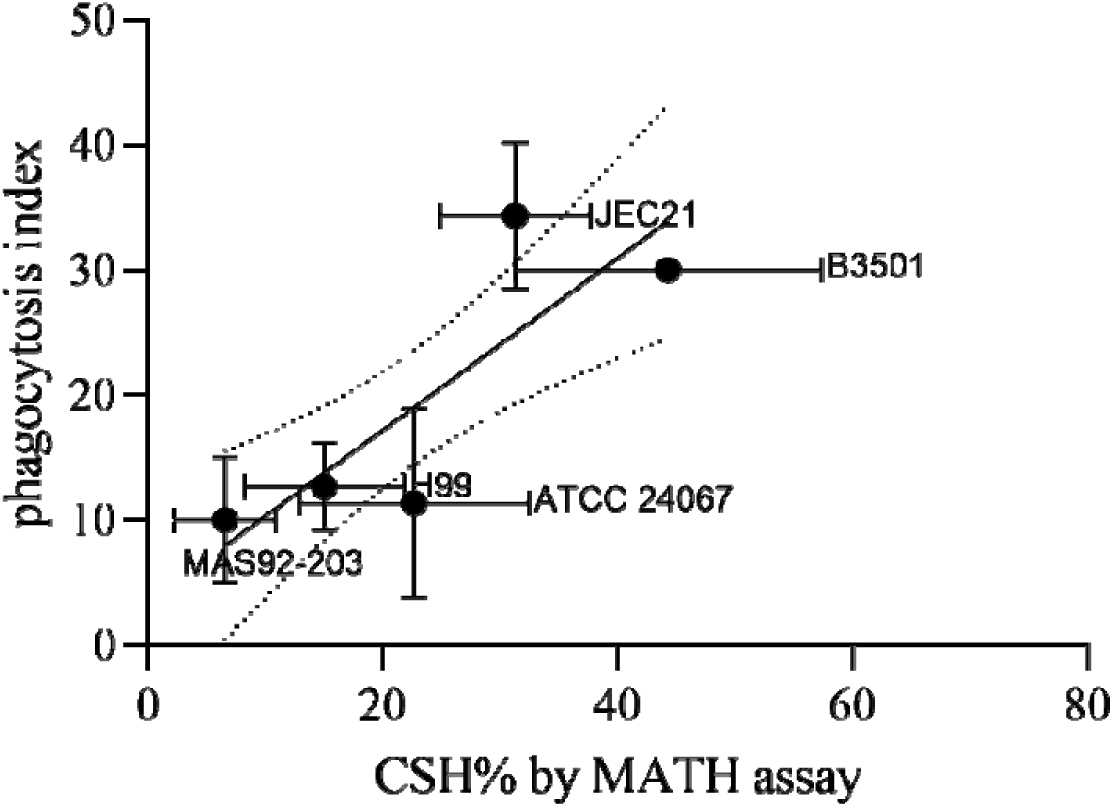
CSH of *C. neoformans* correlates with phagocytosis of *C. neoformans* by natural predator *A. castellani*. Significant positive linear correlation (R^2^ = 0.5722) between CSH of *C. neoformans* strains and phagocytosis index by *A. castellanii*. Phagocytosis index is estimated by fluorescence microscopy as the number of *C. neoformans* labeled by Uvitex internalized per 100 *A. castellanii*. Error bar represents the standard deviation of the mean.

### Effect of antibody binding on CSH

Previous studies have demonstrated that capsule antibody binding alters capsule structure and changes the surface charge of *C. neoformans* (14, 15). This led us to investigate the effect of binding of capsular antibodies to *C. neoformans* on the CSH. We demonstrated that binding of capsular antibody 18B7 (26) increases CSH in a concentration-dependent manner, while binding of non-protective antibody 13F1 has no significant effect on the CSH of *C. neoformans* cells grown in the capsule induction medium (figure 5).

**Figure 5:**
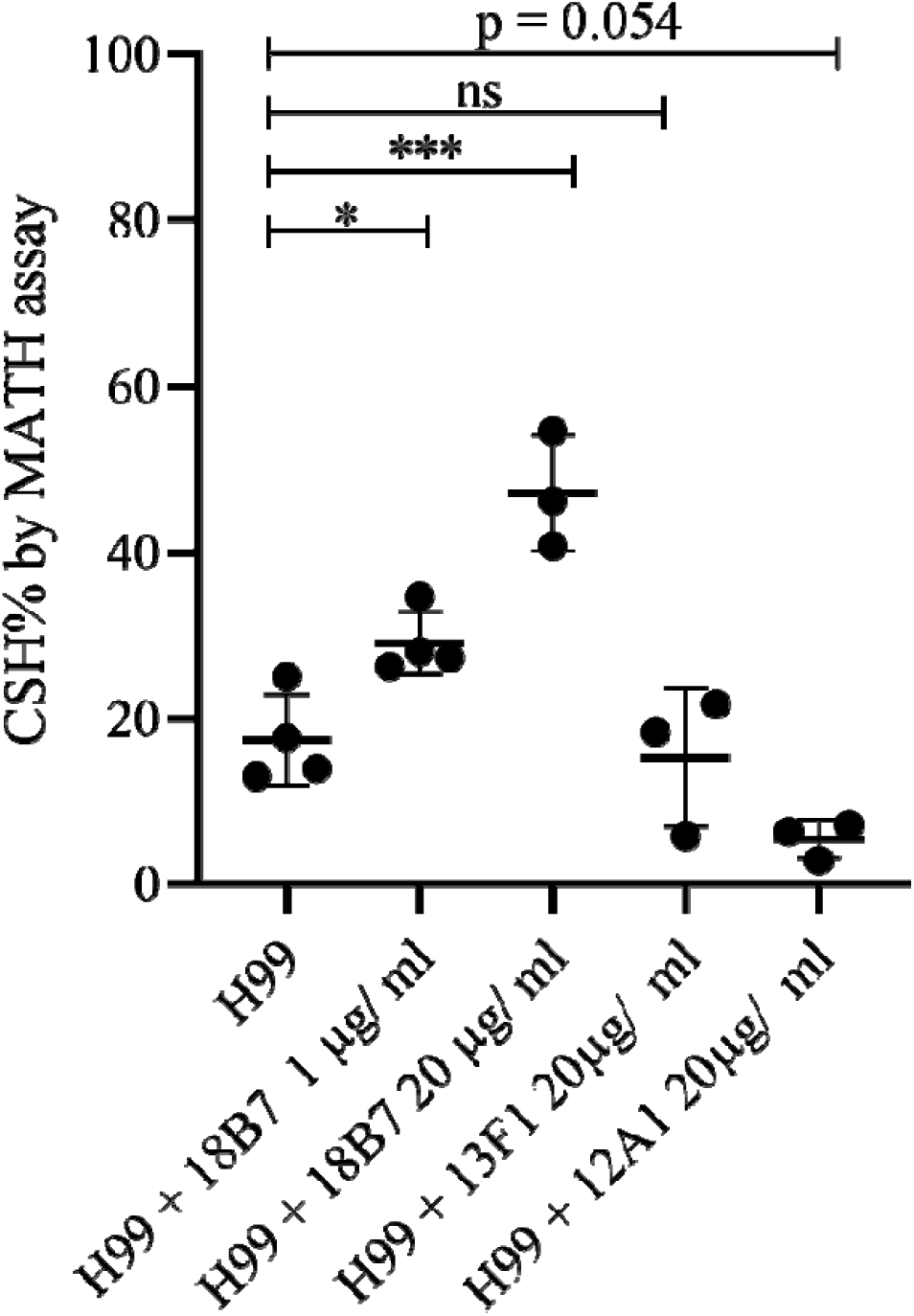
Binding of protective capsule antibodies influences CSH. Incubation of *C. neoformans* strain H99 grown in the capsule induction medium (MM) with protective capsular antibodies 18B7 significantly increase CSH in a concentration-dependent manner, while 12A1 decreased CSH and 13F1 had no significant effect on CSH. CSH was determined by MATH assay in 2-3 biological replicates, as indicated by data points. Error bar represents the standard deviation about the mean. Ordinary one-way ANOVA was used to compare the CSH of untreated *C. neoformans* strain H99 with the CSH of H99 cells treated with different antibodies. The following symbols were used to annotate the statistical significance of the results: ns, p>□0.05; *, p□≤□0.05; **, p□≤□0.01; ***, p□≤□0.001; ****, p□≤□0.0001.

## DISCUSSION

In this study, we measured the CSH of *C. neoformans* and found considerable inter-strain variation. When CSH was estimated by hydrophobic microsphere assay, *C. neoformans* serotype D strains were likely to be more hydrophobic than *C. neoformans* serotype A strain, with the caveat that we analyzed a relatively small set of strains from each serotype. We also demonstrated that CSH is a biophysical parameter that may influence the interaction of yeast cells with the environmental predator *Acanthamoeba castellani*. Finally, we demonstrated that the binding of a protective capsular antibody alters the CSH.

An earlier study suggested that capsule and protective anti-sera binding influenced hydrophobicity of *C. neoformans* (27). They reported no correlation between CSH and phagocytosis of *C. neoformans* by mouse peritoneal macrophages (27). The difference between our observations and the prior report may be attributed to the differences in methodologies. In the prior study, hydrophobicity was estimated from the number of cells that bound to hydrophobic columns. The cells were fixed with formalin, which may have altered surface properties of the yeast. In this study, we have used MATH assay that relies on the interaction of microbe with hydrophobic solvents to calculate CSH (figure 1A) (28). In addition, we have used hydrophobic microsphere assay that quantitates the interaction between hydrophobic beads and the yeast, visualized under a bright field microscope, to estimate the CSH (figure 1B) (18).

*C. neoformans* polysaccharide, like GXM, are essential components required in the formation of microbial communities called biofilms that are protective for the fungi (29). *C. neoformans* biofilms have been reported on medical devices (30, 31). Biofilm associated cells have been associated with increased tolerance against antifungal drugs and phagocytic cells, as they upregulate proteins associated with host defense (32–34). In-vivo, *C. neoformans* form biofilm-like structures called cryptococcomas that could play a role in its neurotropism (35). The surface property of cells may affect the aggregation of microbial communities in biofilms. Interestingly, ATCC24067 and B3501 strains, that are highly hydrophobic also form biofilm more easily when compared to H99 strain that is relatively less hydrophobic (figure 1) (32, 33). A similar correlation between the formation of biofilm and CSH was observed a in *Candida* spp. (19, 36, 37) and in bacteria (38). Flocculation, another multicellular phenotype observed in yeasts, has been observed in *C. neoformans* cells during growth in certain medium (39), and could be caused by changes in CSH, as reported for brewer’s yeast (40).

Amoebas are natural predators of *Cryptococcus* species (20, 41) and have emerged as a powerful tool for studying mechanism of intracellular pathogenesis and evolution of virulence (42, 43). A growing body of evidence suggests that virulence traits have emerged in environmental fungi, including *Cryptococcus* species, because of the selection pressure that results from fungi-amoeba interaction (44). Our finding that the more hydrophobic cryptococcal strains were more readily phagocytosed is congruent with the observation that Amoeba can phagocytose hydrophobic particles (22), although these mechanisms are not well understood. There is a remarkable correspondence between *C. neoformans* virulence traits that influence phagocytosis and enable survival of the fungi in *A. castellanii* and in human macrophages (42). For instance, the capsule of *C. neoformans* masks cell wall components that are recognized by innate immune receptors (45), and the absence of capsule leads to poor survival of *C. neoformans* incubated with *A. castellanii* (42). In-vitro studies of macrophage and *C. neoformans* interaction usually require opsonins such as capsular antibodies, and complement (46, 47) for phagocytosis by innate immune cells. As a result, what is known about the immune response and phagocytosis of *C. neoformans* is greatly influenced by our understanding of host cell receptor-opsonization agent interactions. Studying the effect of CSH on phagocytosis in amoeba may give insights into factors independent of opsonin-receptor interaction, which may influence phagocytosis in macrophages.

Murine antibodies that recognize capsular epitopes of *C. neoformans* can confer passive protection to the host and enhance macrophage activity (48, 49). In addition to facilitating phagocytosis of the yeast, the murine IgG antibody 18B7 (26) alters capsule stiffness and impairs cellular replication of the yeast (15), significantly alters the cell surface charge (14) and has a catalytic activity that breaks down the capsule (50). In this study, we report that mAb 18B7 binding significantly increases the hydrophobicity of *Cryptococcal* cell surface in a concentration-dependent manner, while non-protective antibody, IgM 13F1, did not alter the CSH. We may attribute the differential effect of changes in CSH induced by mAb 13F1 and 18B7 to the pattern of mAb binding, since mAb18B7 binds near the surface in an annular pattern (15, 26, 51), while mAb 13F1 binds throughout the capsule in a punctate pattern (52, 53). There is precedence for our observation in the encapsulated bacteria, *Klebsiella aerogenes*, where the pattern of diffusion of some mAbs through the polysaccharide capsule has been shown to influence the cell surface hydrophobicity (54, 55).

A surprising result in our study was that some *C. neoformans* strains manifest a considerably higher CSH relative to others, despite being surrounded by a hydrophilic capsule. The origin and mechanism for variability in CSH in these strains is not understood. Glycans are intrinsically hydrophilic molecules. Lipophilicity for glycans may be described by the partition coefficient (*P*), that is quantified as the distribution of a compound between two immiscible solvents, like water and octanol (56). While prior studies have compared the lipophilicity for monosaccharides, these efforts are not standardized in the field (13). For small molecules, log *P* can be accurately predicted by an equation proposed by Mannhold *et al.,* although the accuracy of the prediction decreases with an increase in non-hydrogen atoms (25). In this study we used this calculation to predict and compare the lipophilicity of capsular carbohydrate motifs (25), with the caveat that the suitability of these equations for molecules larger than monosaccharides is uncertain. The predicted calculated lipophilicity of GXM oligosaccharides motifs was positive, suggesting that the polymers would preferentially partition into an organic solvent. The M1 motif, which is dominant on the *C. neoformans* serotype D strains, was found to be less lipophilic in comparison to M3 and M4 motifs that are dominant in *C. gattii* serotype B and *C. neoformans* serotype A strains respectively (figure 3). This goes against our experimental observation that some *C. neoformans* serotype D strains were more hydrophobic than serotype B and A strains (figure 2) and implies that simple calculations of lipophilicity do not explain our findings. Instead, we suspect that discrepancy stems from higher-order polysaccharide structures that could present different molecular surfaces in their interaction with the solvent.

The dynamic nature of polysaccharides makes it challenging to obtain defined structures, and to relate the structure of glycans with their activity and biological roles. Yet we know that the flexibility of the oligosaccharide polymer is influenced by intra- and intermolecular hydrogen bonds. Theoretical predictions suggest that α-(1→3)-mannan form weak intermolecular hydrogen bonds, resulting in a polymer with a flexible structure, that allows for many hydroxyl groups to interact with water (11). The primary component in the capsule of *C. neoformans*, is built upon repeating α-(1→3)-mannose triads, which would contribute to the observation that 95% of capsule’s weight comes from water (10). We also found that the number of hydroxyl groups in each motif (figure 3), was inversely related to the observed CSH. The dominant motif M1 in the capsule of *C. neoformans* serotype D had fewer hydroxyl groups and the strains of serotype D tend to have higher CSH, when compared to the number hydroxyl group dominant motifs M2 and M3 of serotype A and B, whose strains had comparatively lower CSH. Fewer hydroxyl groups result in fewer opportunities for hydrogen bonding between the polysaccharide and water, which could translate into less hydrophilic structures with higher CSH.

It is also important to note that the motifs that enrich the capsule may differ between strains of the same serotype (9). For example, *C. neoformans* serotype D strain 24067 a capsular polysaccharide chemotyping suggests that M1 motif dominates 100% of the strain, while *C. neoformans* serotype D strain B3502 is composed of the dominant M1 (52%) motif, and M6 (48%) motif (9). This may contribute to the variation of CSH within strains grouped in serotype D (figure 2, Table S1).

Lipophilic structures have been reported in the capsule, which might extend to the surface and influence the hydrophobicity of the cell surface (11, 57). In addition, the composition of the cell wall, in particular the chitin-chitosan content in the cell wall, that is regulated by CDA genes (58), may influence the hydrophobicity and adhesion of the yeast to various surface (59), a phenomenon that has also been reported in the plant pathogenic fungi *Magnoporthe orzaye* (60). The chemical structures responsible for the high CSH of some strains presents new puzzle for future study.

In summary, we report that CSH of *Cryptococcus* species can differ significantly depending on the strain. We have also demonstrated the correlation of the biophysical parameter CSH, with the phagocytosis by *A. castellanii* and that protective antibodies that bind to the capsule of *C. neoformans* may influence the hydrophobicity of *C. neoformans*. The finding that *C. neoformans* strain differ in CSH, and that changes to this cell-surface property correlates with biological properties, suggests the investigation of how this parameter is established and maintained could provide new insights into capsular structure.

## MATERIALS AND METHODS

### Strains and culture of *C. neoformans* and *C. gattii*

*Cryptococcus neoformans* and *gattii* strains (Table 1) stored as frozen stocks at −80°C were streaked onto Sabouraud Agar plates and incubated at 30°C for 48 hours. The plates were stored at 4°C for use up to 1 week. Multiple colonies were selected and inoculated into 5 ml of liquid media, Sabouraud broth and incubated at 30°C with shaking. For capsule induction, 10^6^ cells/ ml were washed 2X in PBS and inoculated into MM (10□mM MgSO4, 29.3□mM KH_2_PO_4_, 13□mM glycine, 3□µM thiamine-HCl, and 15□mM dextrose, with pH adjusted to 5.5).

**Table 1:** Strains of *C. neoformans* and *C. gattii* used in the present study. The references indicate the study in which the strains were serotyped, or the study in which the strains used had been characterized by serotype.

| Species |  | Mutant | Serotype | Source |
| --- | --- | --- | --- | --- |
| *Cryptococcus neoformans* | H99 |  | A | John Perfect (Durham, NC) |
|  |  | *CAP59* |  |  |
|  | J45 |  |  | (63) |
|  | J10 |  |  | (63) |
|  | J43 |  |  | (63) |
|  | J48 |  |  | (63) |
|  | SB6 |  |  | (64) |
|  | MAS92-203 |  | AD | (6) |
|  | ATCC24067 |  | D | (6) |
|  | B3501 |  |  | (6) |
|  |  | *CAP67* |  |  |
|  | JEC21 |  |  | (65) |
|  | J39 |  |  | (63) |
| *Cryptococcus gattii* | R265 |  | B | ATCC (Manassas, VA) (66) |
|  | NIH444 |  |  | (6) |
|  | WM179 |  |  | ATCC (Manassas, VA)(66) |

### Antibody incubation

*C. neoformans* (H99) grown in MM were washed 2X in PBS. Protective and non-protective capsular antibodies, 18B7, 12A1, and 13F1 (61) respectively, were incubated for 1 hour at 30°C shaking. The CSH% was determined by MATH and microsphere assays, as detailed below.

### Estimation of CSH by MATH

CSH was estimated by MATH assay described in (28). Yeast cultures were washed 2X in PBS and resuspended in 3 mL of PBS at an estimated Initial OD 0.2-0.4 recorded as A_0_. 0.4 ml of n-Hexadecane was added, and the mixture was vortexed for 30 s and incubated at 30°C to allow the layers to separate. Final OD (A_1_) of the aqueous layer was recorded estimated as an average of 3 technical replicates in a 96 well plate read by EMax Plus Microplate Reader (Molecular Devices). CSH% was estimated as [1 – (A_0_/A_1_)] × 100.

### Estimation of CSH by hydrophobic microsphere assay

CSH of *C. neoformans* and *C. gattii* were estimated method detailed in (18) by resuspending 100 µL of 2 × 10^6^ cells/ ml with 9.02 × 10^8^ 0.8 µm green hydrophobic beads (Bang Laboratories) in 2 mL of sodium phosphate buffer (0.05 M, pH 7.2) in clean glass tubes. After equilibration at RT for 2 minutes, the mixture was vortexed vigorously for 30 s. One hundred cells were counted, and the percentage of cells having >3 attached microspheres was considered as the CSH% value.

### *Acanthamoeba castellanii* culture and phagocytosis index

*Acanthamoeba castellanii* strain 30234 was obtained from the American Type Culture Collection (ATCC). Cultures were maintained in PYG broth (ATCC medium 712) at 25°C according to instructions from ATCC.

### *Acanthamoeba castelanii* phagocytosis index

The phagocytosis index was estimated as detailed in (62) with minor modifications. Briefly, 5 × 10^5^ cells/ ml cells of *A. castellanii* were incubated in 35 mm No. 1.5 coverslip MatTek dishes with DPBS (Ca^2+^ and Mg^2+^) for 3-4 hours. *C. neoformans* or *C. gattii* strains were incubated with 10 µg/mL Uvitex (fungal cell wall dye) and inoculated at MOI 1 and incubated for 2 hours at 25°C. The cells were imaged using Zeiss Axiovert 200M inverted microscope with 20× phase objective. Phagocytosis index was estimated by counting the number of *C. neoformans* or *C. gattii* engulfed per 100 amoeboid cells.

### Estimation of lipophilicity and number of hydroxyl groups in carbohydrate motifs

Lipophilicity of the carbohydrate motif dominant in the capsule of *C. neoformans* serotype was estimated by method described by Mannhold *et al*. (25), as the log of the partition coefficient (*P*).

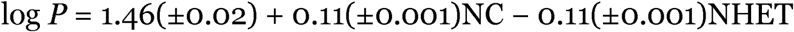

Where NC is the number of carbon atoms in a molecule and NHET is the number of hetero atoms.

The number of hydroxyl groups in each motif of C. neoformans capsule was counted manually, as proxy for the number of hydrogen bond donor and acceptor atoms.

## ACKNOWLEDGMENTS

A.C. was supported by grant 5R01HL059842. C.J.C. was funded by Irish Research Council postgraduate award (GOIPG/2016/998)

R.V. designed and conducted the experiments, analyzed the data, and wrote the manuscript. C.J.C. performed computation and theoretical analysis and wrote the manuscript. A.C. contributed to the experimental design, supervised the experiments, and edited and wrote parts of the manuscript. Special thanks to Radames JB Cordero for valuable discussions of experimental design and edits to the manuscript, and to Daniel Quinn Smith for the valuable contribution of editing the figures.

## SUPPLEMENTARY MATERIAL

**Supplementary Table S1:** Table summarizing the results of multiple comparison of CSH% by MATH assay by ordinary one-way ANOVA, where mean of each column was compared to the mean of the other, (graphical representation in figure 2A).

| Tukey's multiple comparisons test | Mean Diff. | 95.00% CI of diff. | Significant? | Summary | Adjusted P Value |
| --- | --- | --- | --- | --- | --- |
| B3501 vs. H99 | 29.28 | 13.80 to 44.76 | Yes | **** | <0.0001 |
| B3501 vs. ATCC 24067 | 21.62 | 6.141 to 37.10 | Yes | *** | 0.0008 |
| B3501 vs. MAS92-203 | 37.74 | 21.62 to 53.85 | Yes | **** | <0.0001 |
| B3501 vs. R265 | 32.13 | 12.14 to 52.11 | Yes | **** | <0.0001 |
| B3501 vs. WM179 | 27.53 | 9.382 to 45.69 | Yes | *** | 0.0002 |
| B3501 vs. NIH444 | 26.13 | 7.975 to 44.28 | Yes | *** | 0.0005 |
| B3501 vs. J43 | 35.15 | 15.17 to 55.13 | Yes | **** | <0.0001 |
| B3501 vs. SB6 | 26.12 | 2.905 to 49.34 | Yes | * | 0.0149 |
| B3501 vs. JEC21 | 13.01 | -6.970 to 33.00 | No | ns | 0.5717 |
| B3501 vs. J45 | 31.36 | 8.136 to 54.57 | Yes | ** | 0.0013 |
| B3501 vs. J48 | 35.57 | 15.58 to 55.55 | Yes | **** | <0.0001 |
| B3501 vs. J39 | 39.49 | 19.51 to 59.47 | Yes | **** | <0.0001 |
| B3501 vs. CAP59 | 24.32 | 4.335 to 44.30 | Yes | ** | 0.0057 |
| B3501 vs. CAP67 | 23.73 | 3.743 to 43.71 | Yes | ** | 0.0078 |
| H99 vs. ATCC 24067 | -7.659 | -23.14 to 7.820 | No | ns | 0.896 |
| H99 vs. MAS92-203 | 8.456 | -7.656 to 24.57 | No | ns | 0.8496 |
| H99 vs. R265 | 2.846 | -17.14 to 22.83 | No | ns | >0.9999 |
| H99 vs. WM179 | -1.747 | -19.90 to 16.40 | No | ns | >0.9999 |
| H99 vs. NIH444 | -3.154 | -21.31 to 15.00 | No | ns | >0.9999 |
| H99 vs. J43 | 5.87 | -14.11 to 25.85 | No | ns | 0.999 |
| H99 vs. SB6 | -3.156 | -26.38 to 20.06 | No | ns | >0.9999 |
| H99 vs. JEC21 | -16.27 | -36.25 to 3.718 | No | ns | 0.227 |
| H99 vs. J45 | 2.075 | -21.14 to 25.29 | No | ns | >0.9999 |
| H99 vs. J48 | 6.285 | -13.70 to 26.27 | No | ns | 0.9979 |
| H99 vs. J39 | 10.21 | -9.774 to 30.19 | No | ns | 0.8723 |
| H99 vs. *CAP59* | -4.961 | -24.95 to 15.02 | No | ns | 0.9998 |
| H99 vs. *CAP67* | -5.553 | -25.54 to 14.43 | No | ns | 0.9994 |
| ATCC 24067 vs. MAS92-203 | 16.11 | 0.003366 to 32.23 | Yes | * | 0.0499 |
| ATCC 24067 vs. R265 | 10.51 | -9.479 to 30.49 | No | ns | 0.8481 |
| ATCC 24067 vs. WM179 | 5.913 | -12.24 to 24.06 | No | ns | 0.9969 |
| ATCC 24067 vs. NIH444 | 4.505 | -13.65 to 22.66 | No | ns | 0.9998 |
| ATCC 24067 vs. J43 | 13.53 | -6.455 to 33.51 | No | ns | 0.509 |
| ATCC 24067 vs. SB6 | 4.503 | -18.72 to 27.72 | No | ns | >0.9999 |
| ATCC 24067 vs. JEC21 | -8.607 | -28.59 to 11.38 | No | ns | 0.9619 |
| ATCC 24067 vs. J45 | 9.734 | -13.48 to 32.95 | No | ns | 0.9693 |
| ATCC 24067 vs. J48 | 13.94 | -6.040 to 33.93 | No | ns | 0.4594 |
| ATCC 24067 vs. J39 | 17.87 | -2.114 to 37.85 | No | ns | 0.1243 |
| ATCC 24067 vs. *CAP59* | 2.698 | -17.29 to 22.68 | No | ns | >0.9999 |
| ATCC 24067 vs. *CAP67* | 2.106 | -17.88 to 22.09 | No | ns | >0.9999 |
| MAS92-203 vs. R265 | -5.61 | -26.09 to 14.87 | No | ns | 0.9995 |
| MAS92-203 vs. WM179 | -10.2 | -28.90 to 8.491 | No | ns | 0.8115 |
| MAS92-203 vs. NIH444 | -11.61 | -30.30 to 7.084 | No | ns | 0.6453 |
| MAS92-203 vs. J43 | -2.586 | -23.06 to 17.89 | No | ns | >0.9999 |
| MAS92-203 vs. SB6 | -11.61 | -35.26 to 12.03 | No | ns | 0.9011 |
| MAS92-203 vs. JEC21 | -24.72 | -45.20 to -4.244 | Yes | ** | 0.0063 |
| MAS92-203 vs. J45 | -6.381 | -30.03 to 17.26 | No | ns | 0.9996 |
| MAS92-203 vs. J48 | -2.171 | -22.65 to 18.31 | No | ns | >0.9999 |
| MAS92-203 vs. J39 | 1.755 | -18.72 to 22.23 | No | ns | >0.9999 |
| MAS92-203 vs. CAP59 | -13.42 | -33.89 to 7.061 | No | ns | 0.5621 |
| MAS92-203 vs. CAP67 | -14.01 | -34.49 to 6.469 | No | ns | 0.4919 |
| R265 vs. WM179 | -4.592 | -26.71 to 17.53 | No | ns | >0.9999 |
| R265 vs. NIH444 | -6 | -28.12 to 16.12 | No | ns | 0.9996 |
| R265 vs. J43 | 3.024 | -20.62 to 26.67 | No | ns | >0.9999 |
| R265 vs. SB6 | -6.002 | -32.44 to 20.43 | No | ns | >0.9999 |
| R265 vs. JEC21 | -19.11 | -42.76 to 4.534 | No | ns | 0.2361 |
| R265 vs. J45 | -0.7708 | -27.21 to 25.67 | No | ns | >0.9999 |
| R265 vs. J48 | 3.439 | -20.21 to 27.08 | No | ns | >0.9999 |
| R265 vs. J39 | 7.365 | -16.28 to 31.01 | No | ns | 0.9981 |
| R265 vs. CAP59 | -7.807 | -31.45 to 15.84 | No | ns | 0.9965 |
| R265 vs. CAP67 | -8.399 | -32.04 to 15.25 | No | ns | 0.9929 |
| WM179 vs. NIH444 | -1.407 | -21.88 to 19.07 | No | ns | >0.9999 |
| WM179 vs. J43 | 7.616 | -14.50 to 29.73 | No | ns | 0.9947 |
| WM179 vs. SB6 | -1.41 | -26.49 to 23.67 | No | ns | >0.9999 |
| WM179 vs. JEC21 | -14.52 | -36.64 to 7.599 | No | ns | 0.559 |
| WM179 vs. J45 | 3.822 | -21.26 to 28.90 | No | ns | >0.9999 |
| WM179 vs. J48 | 8.031 | -14.09 to 30.15 | No | ns | 0.9912 |
| WM179 vs. J39 | 11.96 | -10.16 to 34.08 | No | ns | 0.8213 |
| WM179 vs. CAP59 | -3.214 | -25.33 to 18.90 | No | ns | >0.9999 |
| WM179 vs. CAP67 | -3.806 | -25.92 to 18.31 | No | ns | >0.9999 |
| NIH444 vs. J43 | 9.024 | -13.09 to 31.14 | No | ns | 0.9755 |
| NIH444 vs. SB6 | -<br>0.00231 | -25.08 to 25.08 | No | ns | >0.9999 |
| NIH444 vs. JEC21 | -13.11 | -35.23 to 9.006 | No | ns | 0.7118 |
| NIH444 vs. J45 | 5.229 | -19.85 to 30.31 | No | ns | >0.9999 |
| NIH444 vs. J48 | 9.439 | -12.68 to 31.56 | No | ns | 0.9646 |
| NIH444 vs. J39 | 13.36 | -8.754 to 35.48 | No | ns | 0.6853 |
| NIH444 vs. *CAP59* | -1.807 | -23.93 to 20.31 | No | ns | >0.9999 |
| NIH444 vs. *CAP67* | -2.399 | -24.52 to 19.72 | No | ns | >0.9999 |
| J43 vs. SB6 | -9.026 | -35.46 to 17.41 | No | ns | 0.9951 |
| J43 vs. JEC21 | -22.14 | -45.78 to 1.510 | No | ns | 0.0879 |
| J43 vs. J45 | -3.795 | -30.23 to 22.64 | No | ns | >0.9999 |
| J43 vs. J48 | 0.4152 | -23.23 to 24.06 | No | ns | >0.9999 |
| J43 vs. J39 | 4.341 | -19.30 to 27.99 | No | ns | >0.9999 |
| J43 vs. *CAP59* | -10.83 | -34.48 to 12.81 | No | ns | 0.9392 |
| J43 vs. *CAP67* | -11.42 | -35.07 to 12.22 | No | ns | 0.9114 |
| SB6 vs. JEC21 | -13.11 | -39.55 to 13.33 | No | ns | 0.8945 |
| SB6 vs. J45 | 5.231 | -23.73 to 34.19 | No | ns | >0.9999 |
| SB6 vs. J48 | 9.441 | -17.00 to 35.88 | No | ns | 0.9925 |
| SB6 vs. J39 | 13.37 | -13.07 to 39.80 | No | ns | 0.8805 |
| SB6 vs. *CAP59* | -1.805 | -28.24 to 24.63 | No | ns | >0.9999 |
| SB6 vs. *CAP67* | -2.397 | -28.83 to 24.04 | No | ns | >0.9999 |
| JEC21 vs. J45 | 18.34 | -8.095 to 44.78 | No | ns | 0.4688 |
| JEC21 vs. J48 | 22.55 | -1.094 to 46.20 | No | ns | 0.0756 |
| JEC21 vs. J39 | 26.48 | 2.831 to 50.12 | Yes | * | 0.0157 |
| JEC21 vs. *CAP59* | 11.31 | -12.34 to 34.95 | No | ns | 0.9175 |
| JEC21 vs. *CAP67* | 10.71 | -12.93 to 34.36 | No | ns | 0.9439 |
| J45 vs. J48 | 4.21 | -22.23 to 30.65 | No | ns | >0.9999 |
| J45 vs. J39 | 8.136 | -18.30 to 34.57 | No | ns | 0.9983 |
| J45 vs. *CAP59* | -7.036 | -33.47 to 19.40 | No | ns | 0.9996 |
| J45 vs. *CAP67* | -7.628 | -34.06 to 18.81 | No | ns | 0.9991 |
| J48 vs. J39 | 3.926 | -19.72 to 27.57 | No | ns | >0.9999 |
| J48 vs. *CAP59* | -11.25 | -34.89 to 12.40 | No | ns | 0.9204 |
| J48 vs. *CAP67* | -11.84 | -35.48 to 11.81 | No | ns | 0.8878 |
| J39 vs. *CAP59* | -15.17 | -38.82 to 8.474 | No | ns | 0.5953 |
| J39 vs. *CAP67* | -15.76 | -39.41 to 7.882 | No | ns | 0.534 |
| *CAP59* vs. *CAP67* | -0.5919 | -24.24 to 23.05 | No | ns | >0.9999 |

**Supplementary Table S2:** Table summarizing the results of multiple comparison of mean CSH% by MATH assay by ordinary one-way ANOVA, where mean of each column was compared to the mean of *C. neoformans* strain H99 (graphically representation in figure 2A).

| Dunnett's multiple comparisons test | Mean Diff. | 95.00% CI of diff. | Significant? | Summary | Adjusted P Value |
| --- | --- | --- | --- | --- | --- |
| H99 vs. B3501 | -29.28 | -42.24 to -16.32 | Yes | **** | <0.0001 |
| H99 vs. ATCC 24067 | -7.659 | -20.62 to 5.298 | No | ns | 0.5766 |
| H99 vs. MAS92-203 | 8.456 | -5.031 to 21.94 | No | ns | 0.4971 |
| H99 vs. R265 | 2.846 | -13.88 to 19.57 | No | ns | 0.9994 |
| H99 vs. WM179 | -1.747 | -16.94 to 13.45 | No | ns | 0.9996 |
| H99 vs. NIH444 | -3.154 | -18.35 to 12.04 | No | ns | 0.9992 |
| H99 vs. J43 | 5.87 | -10.86 to 22.60 | No | ns | 0.9738 |
| H99 vs. SB6 | -3.156 | -22.59 to 16.28 | No | ns | 0.9994 |
| H99 vs. JEC21 | -16.27 | -32.99 to 0.4622 | No | ns | 0.0615 |
| H99 vs. J45 | 2.075 | -17.36 to 21.51 | No | ns | 0.9996 |
| H99 vs. J48 | 6.285 | -10.44 to 23.01 | No | ns | 0.9567 |
| H99 vs. J39 | 10.21 | -6.518 to 26.94 | No | ns | 0.5337 |
| H99 vs. CAP59 | -4.961 | -21.69 to 11.77 | No | ns | 0.9908 |
| H99 vs. CAP67 | -5.553 | -22.28 to 11.18 | No | ns | 0.9837 |

